# Key aspects of neurovascular control mediated by specific populations of inhibitory cortical interneurons

**DOI:** 10.1101/550269

**Authors:** L Lee, L Boorman, E Glendenning, C Christmas, P Sharp, P Redgrave, O Shabir, E Bracci, J Berwick, C Howarth

## Abstract

Inhibitory interneurons can evoke vasodilation and vasoconstriction, making them potential cellular drivers of neurovascular coupling. However, the specific regulatory roles played by particular interneuron subpopulations remain unclear. Our purpose was therefore to adopt a cell-specific optogenetic approach to investigate how somatostatin (SST) and neuronal nitric oxide synthase (NOS1)-expressing interneurons might influence neurovascular relationships. In mice, specific activation of SST- or NOS1-interneurons was sufficient to evoke haemodynamic changes similar to those evoked by physiological whisker stimulation. In the case of NOS1-interneurons, robust haemodynamic changes occurred with minimal changes in neural activity. Conversely, activation of SST-interneurons produced robust changes in evoked neural activity with shallow cortical excitation and pronounced deep layer cortical inhibition. This often resulted in a central increase in blood volume with corresponding surround decrease, analogous to the negative BOLD signal. These results demonstrate the role of specific populations of cortical interneurons in the active control of neurovascular function.

## Introduction

Neurovascular coupling (NVC) is the mechanism through which local cerebral blood flow (CBF) changes are tightly coupled to increases in neural activity. Since the reserves of oxygen and glucose within neurons are strictly limited, such coupling is essential for normal brain function. Variations in blood oxygenation and volume evoked by neural activity underlie functional imaging signals, such as blood oxygen level dependent functional magnetic resonance imaging (BOLD-fMRI), which are commonly used as a surrogate measure of local changes in neuronal activity. While most research has focussed on the ability of excitatory neurons (Lacroix et al., 2015, Lecrux et al., 2011) and astrocytes (Mulligan and MacVicar, 2004, Zonta et al., 2003, Lind et al., 2013) to elicit changes in CBF, there has been less focus on the role of inhibitory neurons. GABAergic interneurons innervate local microvessels (Cauli et al., 2004, Vaucher et al., 2000, Hamel, 2006) and have been shown to induce both vasodilation and constriction (Cauli et al., 2004), making them potential cellular drivers of NVC.

Although recent studies have investigated the contribution of inhibitory interneurons to cerebral blood flow regulation by using an optogenetic approach targeting VGAT-expressing neurons (Uhlirova et al., 2016, Anenberg et al., 2015, Vazquez et al., 2018), the role of specific subpopulations of GABAergic interneurons remains unknown. Somatostatin (SST)-expressing neurons, which account for around 30% of GABAergic interneurons in the somatosensory cortex (Rudy et al., 2011), contact brain microvessels, in particular those in the superficial layers of the cortex (Kocharyan et al., 2008). The release of GABA by SST interneurons has been suggested to contribute to basal forebrain stimulation-evoked cortical CBF responses (Kocharyan et al., 2008). In addition, approximately 28% of GABAergic neurons (Cauli et al., 2004), including a small subset of SST interneurons (Yavorska and Wehr, 2016, Karagiannis et al., 2009), express neuronal nitric oxide synthase (NOS1). This further subpopulation of interneurons releases nitric oxide (NO), which has for a long time been known to be a potent vasodilator (Ignarro et al., 1987, Palmer et al., 1987, Furchgott and Zawadzki, 1980). NOS1 interneurons are therefore of particular interest in terms of CBF regulation. To investigate the role of these two subpopulations of GABAergic interneuron we used a cell-type specific optogenetic approach that specifically targeted SST- or NOS1-expressing interneurons. By separately activating these two subsets of inhibitory interneurons (those expressing SST or NOS1) we sought to determine how they might regulate cortical haemodynamics. We were able to show that activating both subsets of interneurons evoked a localised haemodynamic response. Importantly, in the case of optogenetic-activation of NOS1 interneurons, the observed haemodynamic changes occurred with only a minimal change in measured multiunit neural activity. Alternatively, after activating SST interneurons negative haemodynamic responses were sometimes observed in the cortical areas surrounding the local area of optogenetic stimulation. This observation is similar to reported negative BOLD fMRI responses, which have been linked to inhibitory neuron activity (Shmuel et al., 2006, Boorman et al., 2015, Shmuel et al., 2002, Stefanovic et al., 2004, Boorman et al., 2010). These observations suggest that specific subpopulations of cortical GABAergic interneurons have specific roles in NVC. Also, that the ability of BOLD signals to act as a surrogate measure of local neural activation may in part be dependent upon which subpopulation of neurons are being activated.

## Results

### Short duration optogenetic stimulation of specific interneurons evokes a localised haemodynamic response

Genetically modified mice expressing channelrhodopsin-2 (ChR2) in either SST- or NOS1-expressing interneurons (referred to as SST-ChR2 or NOS1-ChR2 mice, respectively) were used to investigate how light induced activity of these inhibitory interneurons may alter cortical haemodynamics. Using an anaesthetised mouse (Figure 1), we assessed whether short duration optogenetic stimulation of specific subtypes of interneuron evoked a localised haemodynamic response, comparable to that evoked by a mild physiological stimulus (mechanical whisker stimulation). 2-dimensional optical imaging spectroscopy (2D-OIS) was used to record high-resolution 2D maps of the changes in blood volume (Hbt), oxygenated haemoglobin (HbO_2_) and reduced haemoglobin (Hbr) evoked by stimulation. Each animal initially received a mechanical whisker stimulation (2s, 5Hz), evoking changes in Hbt, HbO_2_ and Hbr which were localised to the whisker barrel cortex (Figure 2A). These haemodynamic changes allowed us to map the whisker barrel cortex and, in turn, guide the placement of the optical fibre used for photostimulation (Figure 1). The time series of the haemodynamic response to whisker stimulation shows an increase in Hbt and HbO_2_ during the stimulation with a corresponding washout of Hbr (Figure 2A).

**Figure 1:**
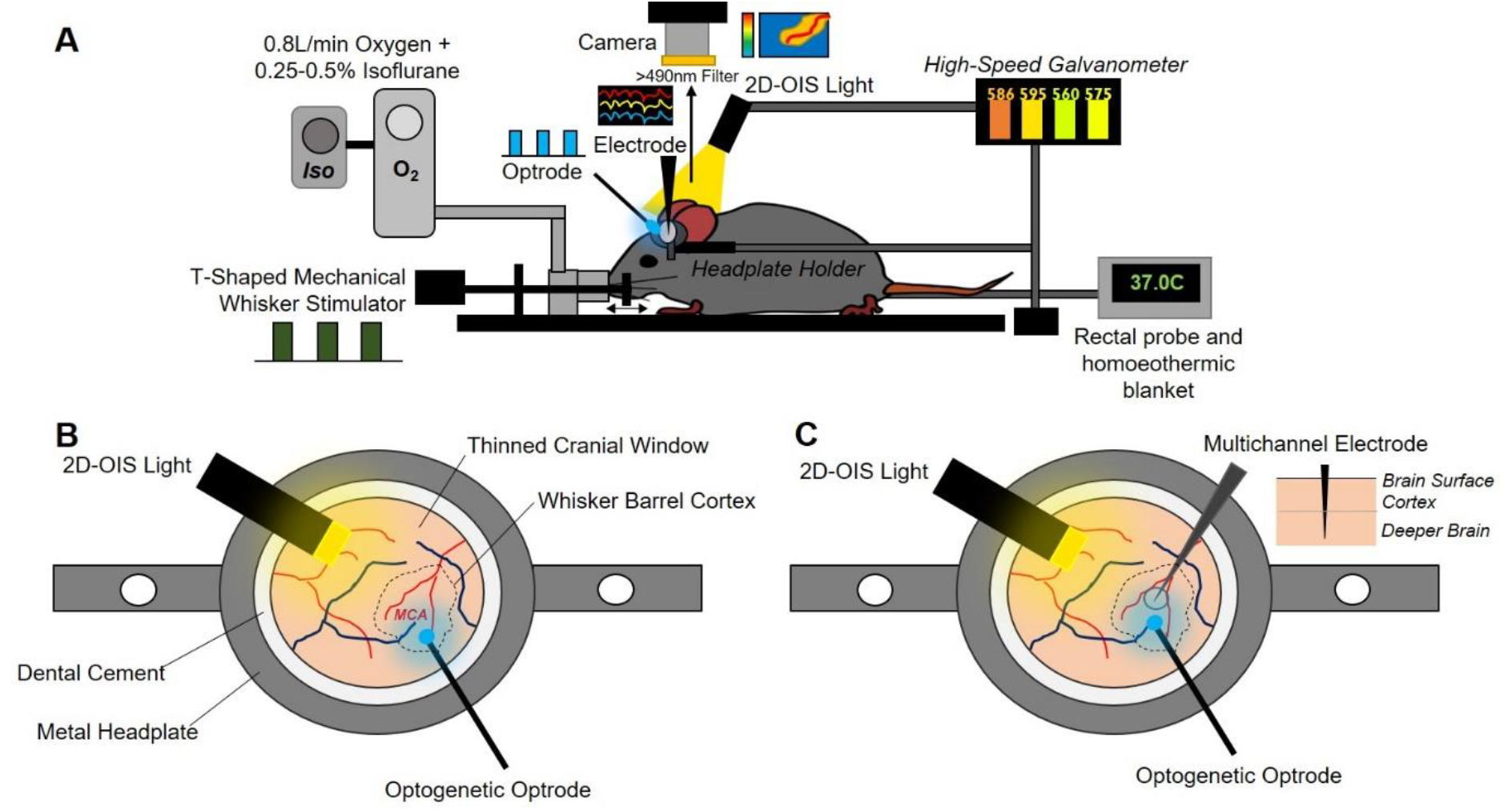
Chronic imaging preparation. **(A)** Imaging setup showing inhalational anaesthetic maintenance, mechanical whisker stimulation, temperature regulation. haemodynamic imaging (2D-OIS). optogenetic stimulation & multichannel electrode electrophysiology **(B)** 1^st^ Imaging session 2-weeks post-surgery with optogenetic optrode placed over MCAI whisker barrel cortex region **(C)** 2^nd^ imaging session 3-weeks post surgery with electrode inserted into whisker barrel cortex + optrode for optogenetic stimulation

**Figure 2:**
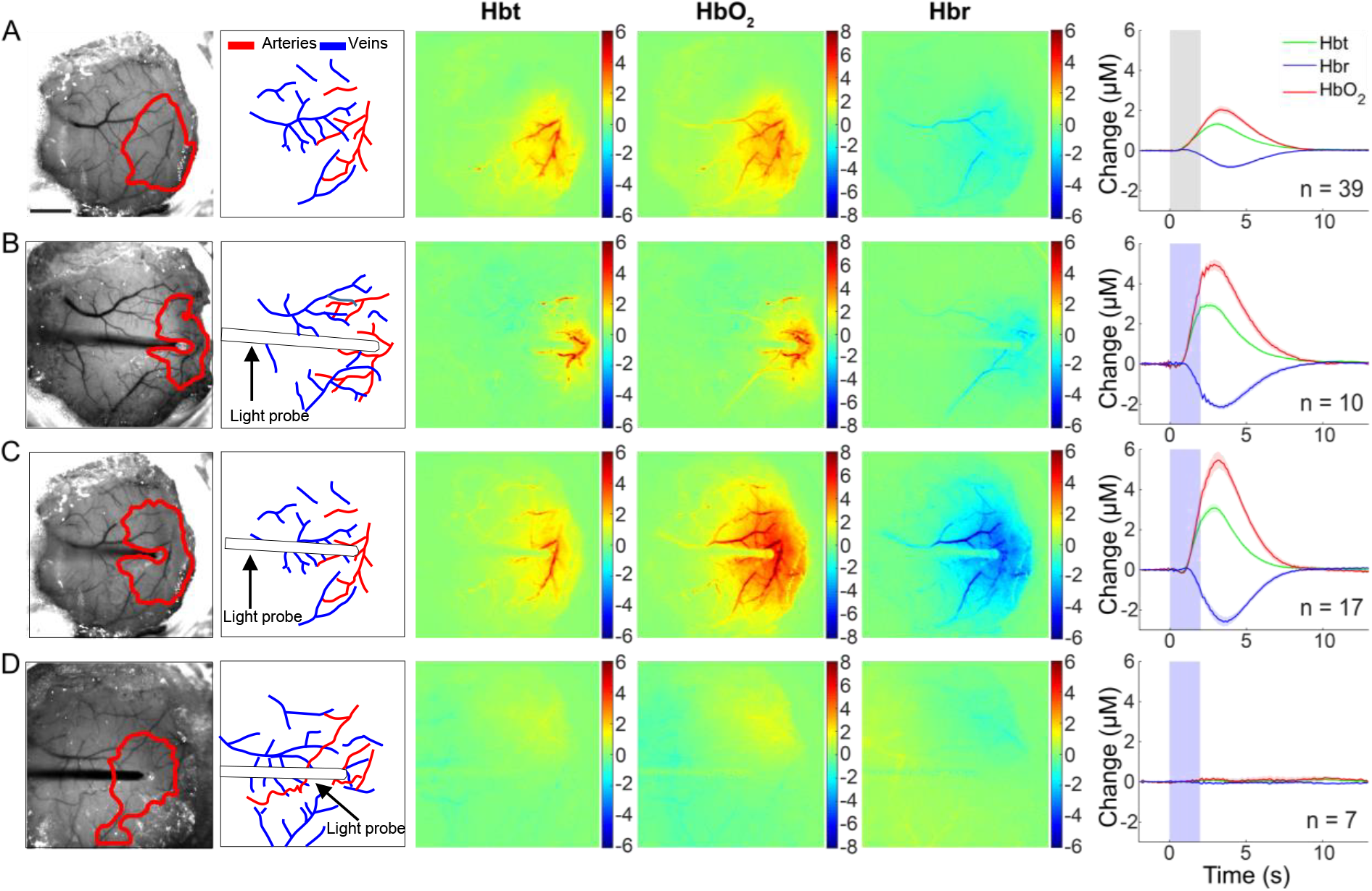
Haemodynamic responses to 2s stimulation. **(A)** mechanical whisker stimulation (5Hz) **(B)** photostimulation (99Hz, 1V or 20Hz 2V) of SST-ChR2 mouse **(C)** photostimulation of NOS1-ChR2 mice **(D)** photostimulation of control mice. 1^st^ column: Representative surface vasculature overlying somatosensory cortex (at 575nm illumination), region of interest (ROI) for analysis shown in red. Scalebar represents 1mm. 2^nd^ column: Vessel map showing surface arteries and veins. 3^rd^, 4^th^ and 5^th^ column: Representative spatial activation map of trial-averaged changes in [Hbt], [HbO_2_] and [Hbr], respectively, with respect to baseline during stimulation. 6^th^ column: Haemodynamic time series within whisker barrel cortex ROI (mean ± s.e.m). Grey box indicates whisker stimulation, blue box indicates photostimulation.

A fibre-coupled blue (470nm) LED, placed directly above the whisker barrel cortex, was used to apply photostimulation in order to activate ChR2-expressing neurons. The fibre optic was positioned in the centre of the whisker barrel cortex (Figure 1) and short duration photostimulation (2s: 99Hz, 1V and 20Hz, 2V) was applied. Similarly to whisker stimulation, for both SST-ChR2 and NOS1-ChR2 mice, a short 2s stimulation produced a robust haemodynamic response focussed around the tip of the fibre optic light guide. Photostimulation elicited a functional hyperemia response, showing a localised increase in blood volume (Hbt) and oxygen saturation (HbO_2_) with corresponding washout of Hbr (Figure 2B,C). These haemodynamic changes were similar in shape to those seen in response to whisker stimulation (Figure 2A) and were within the range of physiological responses (Gu et al., 2018). As was observed with whisker stimulation, increases in Hbt and HbO_2_ were strongest in the middle cerebral artery (MCA) branches overlaying the whisker barrel cortex with a decrease in Hbr which was clearly apparent in the draining veins (Figure 2). When comparing peak values of the evoked Hbt response, a one-way ANOVA showed an overall effect of stimulation type (F_3,69_ = 51.88, p<0.0001) however, Tukey’s multiple comparisons test showed that there was no significant difference in Hbt peak between SST-ChR2 and NOS1-ChR2 mice (p = 0.9531).

Expression of ChR2 in the appropriate cell types was confirmed using immunohistochemistry (Figure 3). Expression of ChR2 was evidenced by the presence of enhanced yellow fluorescent protein (EYFP), the reporter for ChR2 expression, and co-localisation was seen with either SST (Figure 3A) or NOS1 (Figure 3B), as appropriate.

**Figure 3:**
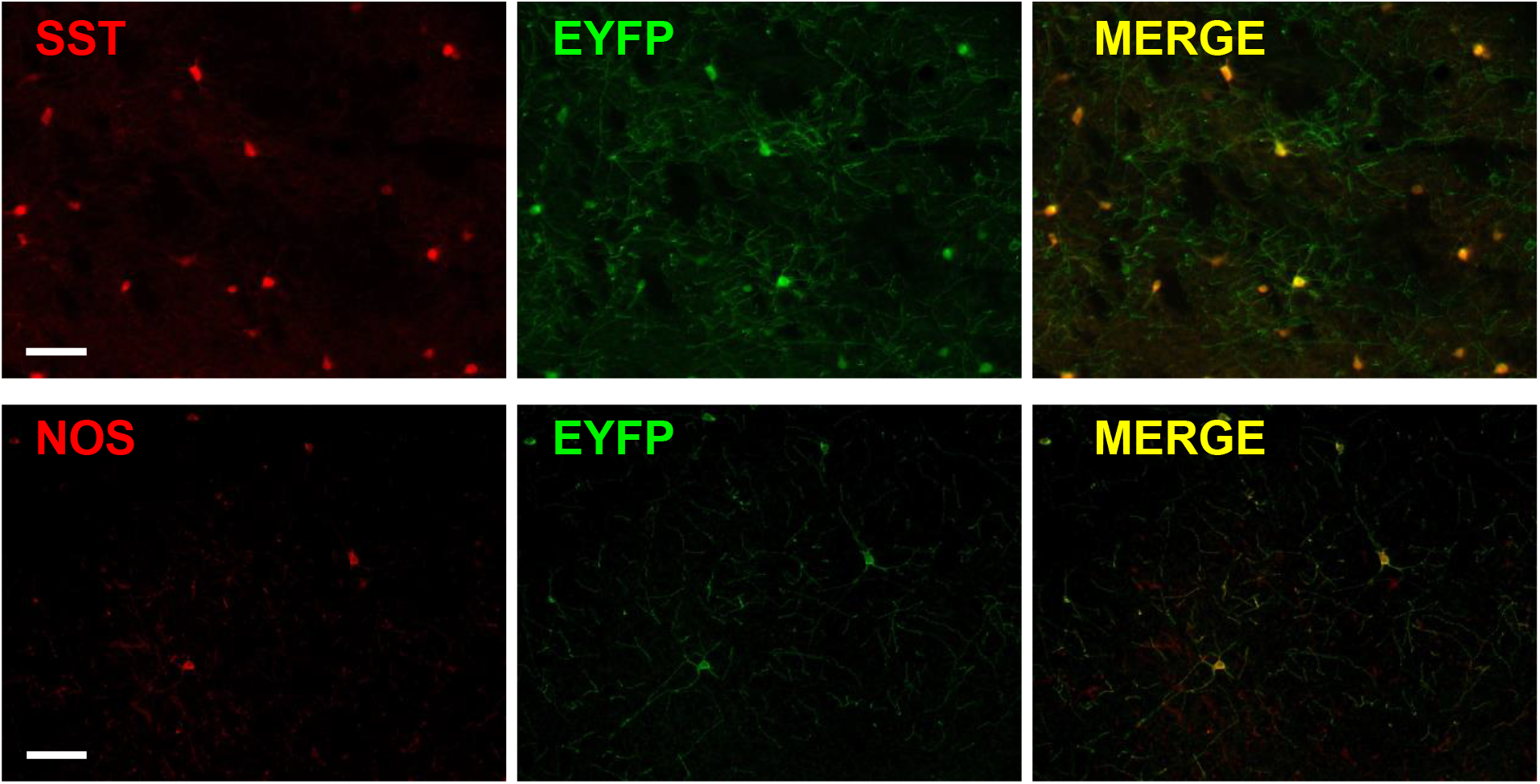
Cell-specific expression of ChR2. Immunohistochemistry showing staining for **(A)** SST (red) and ChR2 (green, EYFP is the reporter for ChR2) and merge (yellow) in cortex from SST-ChR2 mouse. **(B)** NOS1 (red) and ChR2 (green, EYFP is the reporter for ChR2) and merge (yellow) in cortex from NOS1-ChR2 mouse. Scalebar represents 100µm.

Previous studies have reported photostimulation-evoked fMRI (Christie et al., 2013) and cerebral blood flow (Rungta et al., 2017) responses in optogenetically naïve (non-ChR2-expressing) animals. While these studies used a laser to drive the optogenetic response, which may cause issues with heating, our light stimulation used a cold LED light source. To confirm our response was not an artefact we applied our photostimulation paradigm to control animals (either C57bl/6J or non-ChR2-expressing littermates of NOS1-ChR2 animals) and confirmed that such stimulation failed to elicit haemodynamic responses (Figure 2D).

These data demonstrate that short duration activation of specific interneuron subpopulations is sufficient to induce a localised haemodynamic response similar to that evoked by mild mechanical somatosensory stimulation.

### Electrophysiological response to short duration stimulation is dependent on the specific interneuron population activated

Having demonstrated that specific activation of either SST or NOS1 interneurons resulted in localised haemodynamic responses, in a second experimental session we assessed the associated evoked electrophysiological activity. 16 channel Neuronexus probes were inserted into the centre of the whisker barrel cortex in order to measure electrophysiological responses to both whisker stimulation and photostimulation (Figure 1). Electrophysiological and haemodynamic measurements were made concurrently. Across all animals, 2s mechanical whisker stimulation evoked a typical electrophysiological response, with peak local multi-unit activity (MUA) centred around layer 4 of the cortex. The evoked increases in multi-unit activity extended throughout the cortex (Figure 4A).

**Figure 4:**
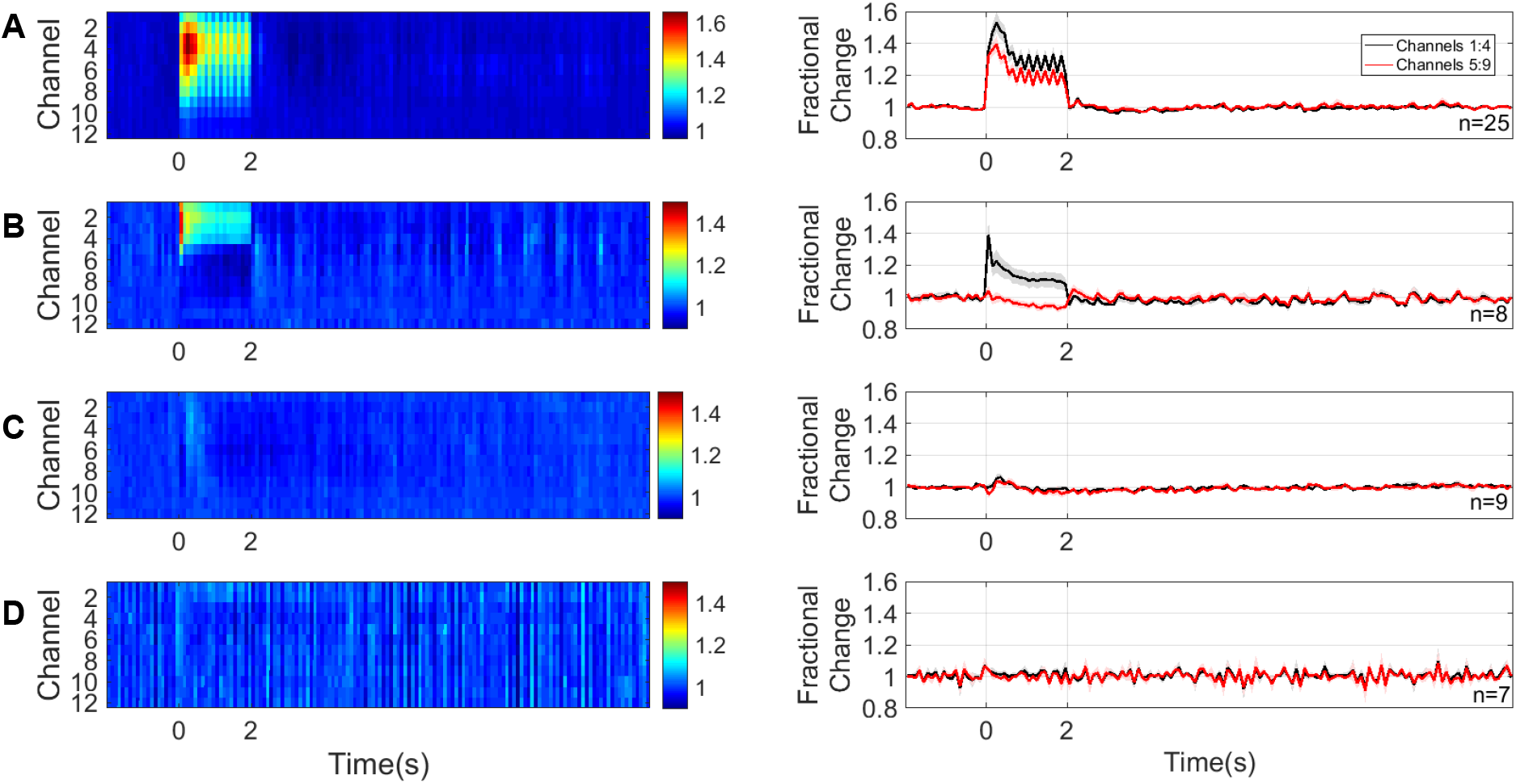
Neural responses to 2s stimulation. **(A)** mechanical whisker stimulation (5Hz) **(B)** photostimulation (99Hz, 1V or 20Hz 2V) of SST-ChR2 mice **(C)** photostimulation of NOS1-ChR2 mice **(D)** photostimulation of control mice. 1^st^ column: Mean MUA response across cortical layers. Colour bar represents fractional change. 2^nd^ column: time series of response through different cortical layers. Data shown are mean ± s.e.m.

Two second photostimulation of SST interneurons evoked an increase in local MUA which was limited to the superficial depth of the cortex (mean fractional change: channels 1:4=1.14±0.05, compared to channels 5:9=0.96±0.02, p = 0.0026, n=8, paired t test, Figure 4B). These data confirm that photostimulation of ChR2-expressing SST interneurons results in measurable changes in neural activity in the superficial depth of the cortex.

Surprisingly, the equivalent photostimulation of ChR2-expressing NOS1 interneurons elicited a minimal change in local neural activity during the light stimulation period (Figure 4C). Given the robust haemodynamic response evoked by activation of NOS1 interneurons (Figure 2C), such a minimal change in neural activity was unexpected.

We confirmed that there were no measurable changes in neural activity evoked by photostimulation in control animals (Figure 4D).

When comparing photostimulation-evoked MUA in channels 1-4 across the groups of animals, a one-way ANOVA showed an overall significant effect (F_2,21_ = 6.223, p =0.0075). Tukey’s multiple comparisons test found that the neural activity evoked in SST-ChR2 mice was significantly different to that evoked in NOS1-ChR2 (p = 0.0092) and control animals (p=0.0325). Taken together, our haemodynamic and electrophysiology data suggest that, in response to short duration activation, both SST and NOS1 interneurons are able to evoke robust changes in blood volume and saturation. However, in the case of specific NOS1 interneuron activation, the robust haemodynamic response occurred with only minimal change in local MUA.

### Long duration stimulation evokes a localised haemodynamic response whose time course differs depending on the specific interneurons activated

Having assessed the responses evoked by short duration interneuron stimulation, in the same group of animals we also performed long duration (16s) stimulation experiments across both modalities (whisker and photostimulation). Mechanical whisker stimulation (16s, 5Hz) was applied and the evoked haemodynamic changes observed using 2D-OIS. Long duration whisker stimulation evoked changes in Hbt, HbO_2_ and Hbr which were localised to the whisker barrel cortex (Figure 5A). The time series of the haemodynamic response to whisker stimulation shows an increase in Hbt and HbO_2_ during the stimulation period with a corresponding washout of Hbr (Figure 5A). The shape of these responses are comparable to those previously reported in awake animals (Sharp et al., 2015).

**Figure 5:**
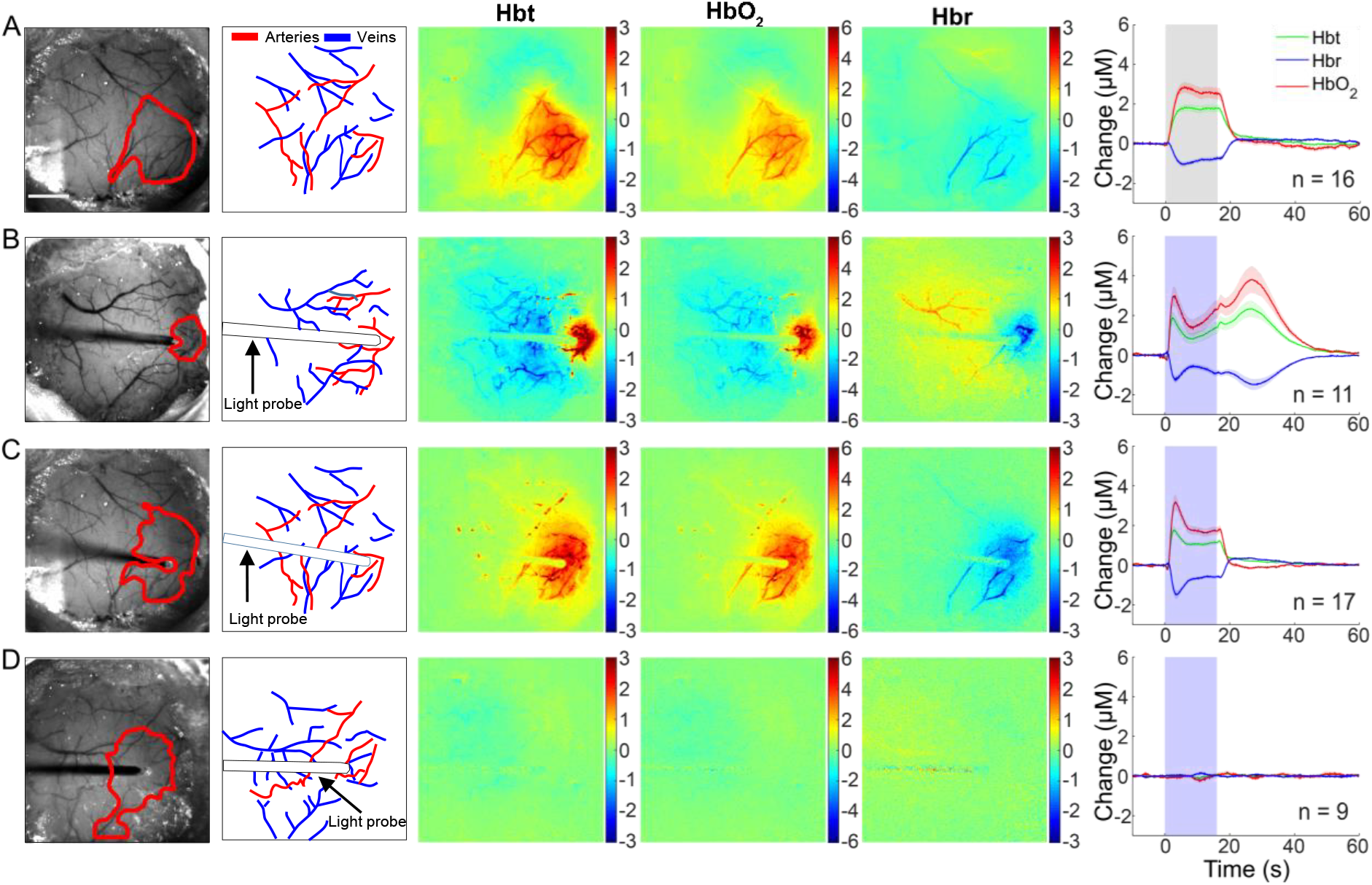
Haemodynamic responses to 16s stimulation. **(A)** mechanical whisker stimulation (5Hz) **(B)** photostimulation (99Hz, 0.5V or 20Hz 1.5V) of SST-ChR2 mouse **(C)** photostimulation of NOS1-ChR2 mice **(D)** photostimulation of control mice. 1^st^ column: Representative surface vasculature overlying somatosensory cortex (at 575nm illumination), region of interest (ROI) for analysis shown in red. Scalebar represents 1mm. 2^nd^ column: Vessel map showing surface arteries and veins. 3^rd^, 4^th^ and 5^th^ Column: Representative spatial activation map of trial-averaged changes in [Hbt], [HbO_2_] and [Hbr], respectively, with respect to baseline during stimulation. 6^th^ column: Haemodynamic time series within whisker barrel cortex ROI (mean ± s.e.m). Grey box indicates whisker stimulation, blue box indicates photostimulation. White arrowheads indicate negative surround response.

LED photostimulation was performed as described above but with the stimulation period extended to 16 seconds. Specific activation of either SST (Figure 5B) or NOS1 (Figure 5C) interneurons resulted in a localised haemodynamic response at the point of stimulation which consisted of an initial increase in Hbt and HbO_2_ with a corresponding washout of Hbr, similar to those elicited by whisker stimulation. 16s photostimulation failed to elicit haemodynamic responses in naïve animals (Figure 5D). When comparing peak evoked haemodynamic responses, a one-way ANOVA showed a significant effect of stimulation type (F_3,49_=12.6, p<0.0001) on Hbt response. However, Tukey’s multiple comparisons test found that this was only significant in the case of photostimulation of naïve animals (Hbt peak: compared to whisker stimulation, p<0.0001; compared to SST activation, p<0.0001; compared to NOS1 activation, p<0.0001).

In the case of photostimulation of SST-ChR2 mice, although the initial haemodynamic changes in the central region were similar to those in response to whisker stimulation, there were some important differences. Some SST-ChR2 animals showed a robust negative surround haemodynamic response (see white arrowheads on representative images in Figure 5B), where the haemodynamic changes were the opposite of the centrally activated region. SST-ChR2 mice also showed a robust increase in blood volume and saturation after the cessation of the 16s optical stimulation, these observations will be described further below. In contrast, no evidence of either a negative surround or secondary haemodynamic response was observed following specific activation of NOS1 interneurons.

### Electrophysiological response to long duration stimulation is dependent on specific interneuron population activated

Having demonstrated that longer duration optogenetic activation of either SST- or NOS1-expressing interneurons reliably results in haemodynamic responses, in a second experimental session we assessed the associated evoked electrophysiological activity. These experiments were performed in the same group of animals as the short duration stimulation investigations. In all animals, 16s mechanical whisker stimulation evoked a typical electrophysiological response which extended throughout the cortex (Figure 6A).

**Figure 6:**
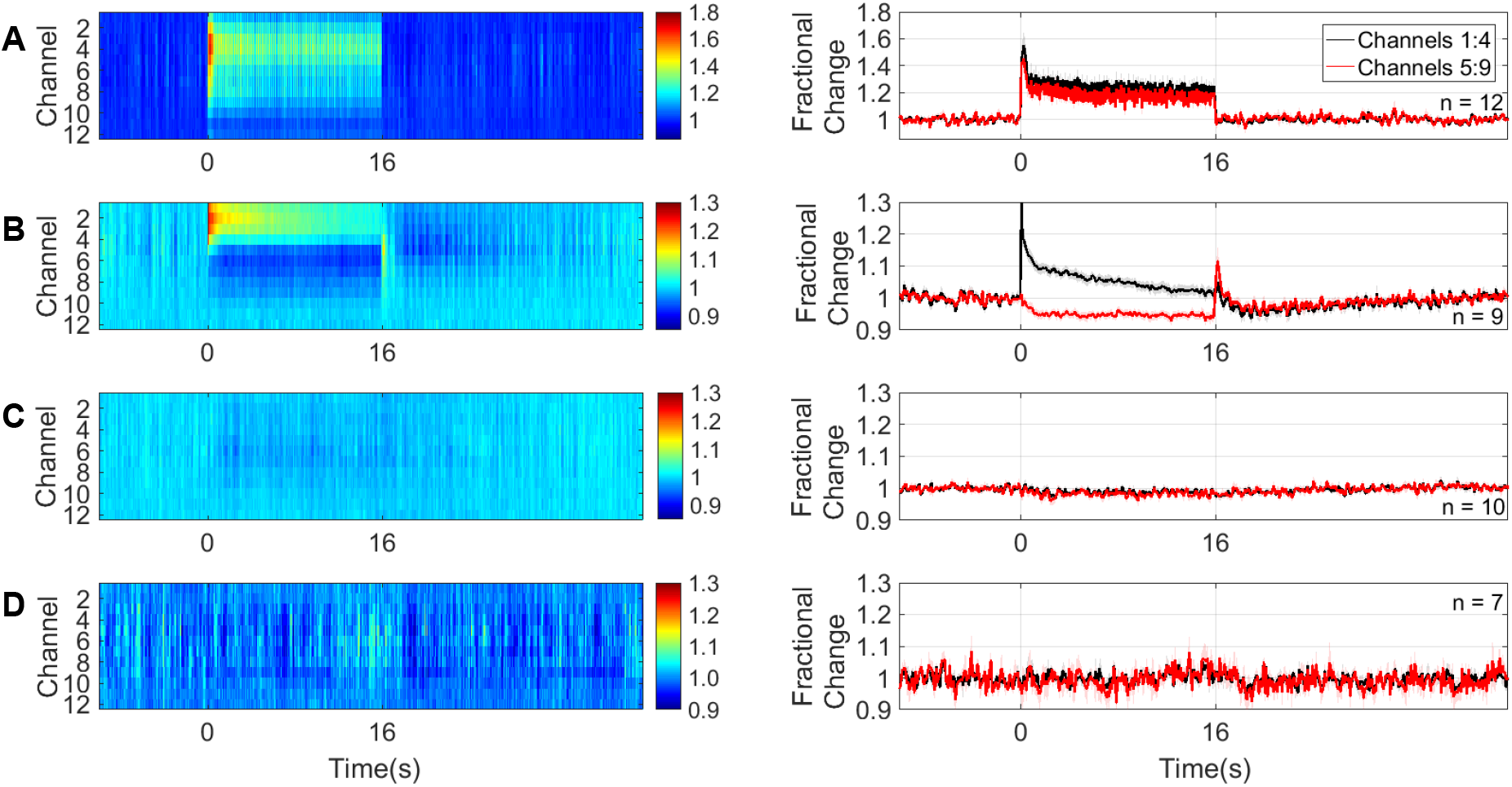
Neural responses to 16s stimulation. **(A)** mechanical whisker stimulation (5Hz) **(B)** photostimulation (99Hz, 0.5V or 20Hz 1.5V) of SST-ChR2 mice **(C)** photostimulation of NOS1-ChR2 mice **(D)** photostimulation of control mice. 1^st^ column: Mean MUA response across cortical layers. Colour bar represents fractional change. 2^nd^ column: time series of response through different cortical layers. Data shown are mean ± s.e.m.

16s photostimulation of SST-expressing interneurons evoked an increase in local MUA in the superficial depth of the cortex and a reduction in MUA deeper in the cortex (mean fractional change in MUA: channels 1:4, 1.06±0.01, n=9, compared to channels 5:9, 0.95±0.01, n=10, p <0.0001, unpaired t test, Figure 6B). Following cessation of the photostimulation there was a brief increase in MUA in the deeper layers, followed by a prolonged decrease below baseline in MUA which extended across cortical depth (0.97±0.01 channels 1:4, 0.98±0.01 channels 5:9, p = 0.6, unpaired t test). These data confirm that specific activation of SST interneurons results in measurable changes in local neural activity in the cortex, the polarity of which is dependent on cortical depth.

An equivalent 16s photostimulation of NOS1-expressing interneurons elicited a minimal change in local neural activity during the light stimulation period (Figure 6C), Although this result is surprising, given the robust haemodynamic response evoked by the photostimulation (Figure 5C), it is consistent with the minimal MUA changes in response to 2s photostimulation of NOS1-expressing interneurons (Figure 4C).

We confirmed that, as expected, no changes in neural activity were detected in response to 16s photostimulation of control animals (Figure 6D).

### Long duration stimulation of SST-expressing interneurons can evoke a negative surround haemodynamic response

When assessing the effect of long duration activation of SST interneurons, in addition to the positive haemodynamic response observed in the central activated region (surrounding the optic fibre), a negative surround haemodynamic response was observed in 9/16 experiments. The negative surround haemodynamic response, which occurs in the cortical region surrounding the central area, consisted of a decrease in Hbt and HbO_2_ and increase in Hbr (white arrowheads, Figure 5B). Interestingly, the temporal dynamics of the blood volume reduction in the central region (Figure 7A, red trace) were almost identical to those seen in the surround (Figure 7A, blue trace). In order to further investigate the origins of the negative surround haemodynamic response, and to assess whether there was a neural marker associated with its presence, experiments in which simultaneous multi-channel electrophysiology and haemodynamic responses were recorded in the central region, were split into those in which a negative surround haemodynamic response was (Figure 7A, n = 9 experiments, 6 mice) and was not (Figure 7B, n = 7 experiments, 6 mice) observed in response to photostimulation. In those experiments in which a negative surround response was observed (decreased Hbt in response to photoactivation of SST neurons, Figure 7A) a second increase in Hbt was also observed. This secondary haemodynamic response, which lasted for around 20 seconds after the photostimulation period ended, was absent in experimental sessions in which there was no negative surround haemodynamic response (Figure 7B). In the central region, both the initial Hbt increase which occurred during light stimulation and the secondary Hbt increase were of similar magnitude (initial peak: 2.12±0.48µM, secondary peak: 2.81±0.57µM, p = 0.39, paired t test, Figure 7A; centre region).

**Figure 7:**
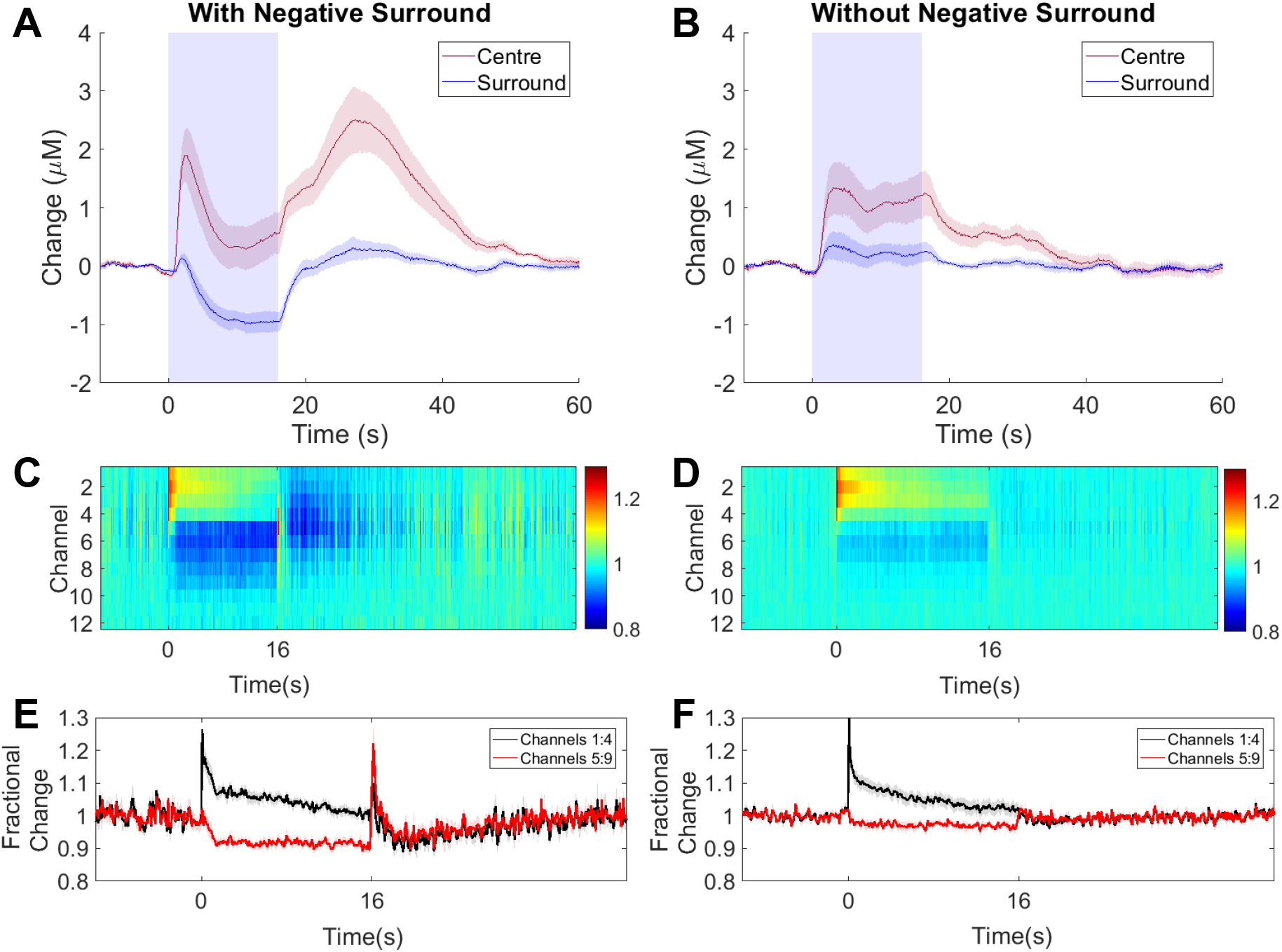
16s photostimulation elicits a negative surround in some SST-ChR2 mice. **(A,B)** Mean time series showing mean changes in [Hbt] with respect to baseline in SST-ChR2 mice which did (**A**, n=9/16 experiments) and did not (**B**, n=7/16 experiments) display a negative surround haemodynamic response. Blue box indicates photostimulation. Response in centre region shown in red, surround region in blue. **(C,D)** Mean MUA response in central region across cortical layers for experiments which did (**C**, n=9 experiments from 6 mice) and did not (**D,** n=7 experiments from 6 mice) display a negative surround haemodynamic response. Colour bar represents fractional change. **(E,F)** Time series of response through different cortical layers for experiments which did (**E**) and did not (**F**) display a negative surround haemodynamic response. Data shown are mean ± s.e.m.

16s photoactivation of SST interneurons evoked an increase in MUA in the superficial depth of the cortex and a decrease in MUA deeper in the cortex. The neuronal response in the superficial layers of the cortex was similar in all experiments, however, the reduction in MUA occurring in the deeper cortical layers was significantly stronger in experiments displaying a negative surround haemodynamic response (Figure 7C-F). A two-way ANOVA showed a significant effect of both electrode channel depth (F_1,26_=59.47, p<0.0001) and presence of negative surround haemodynamic response (F_1,26_=5.204, p = 0.031), but not for their interaction (F_1,26_=3.61, p=0.0686). Tukey’s multiple comparisons test found that the change in MUA in deeper cortical layers (channels 5:9) was significantly different in the presence and absence of a negative surround haemodynamic response (p = 0.0253). Furthermore, in experiments which displayed a negative surround response, at the end of stimulation there was a brief transient increase in MUA in the deeper cortical layers, followed by a long lasting period (20s) of suppressed MUA below baseline which extended throughout the depth of the cortex (Figure 7C,E). Neither of these phenomena were observed in experiments which were lacking a negative surround haemodynamic response (Fig 7D,F).

## Discussion

The present *in vivo* study investigated the contribution of two populations of GABAergic interneurons to neurovascular coupling using complementary optogenetic, imaging (2D-OIS), and electrophysiological procedures. We demonstrated that photoactivation of either SST- or NOS1-expressing interneurons was sufficient to evoke a robust localised haemodynamic response. This shows that these two subsets of interneurons individually are able to drive changes in cerebral haemodynamics. In the case of specific activation of SST interneurons, prolonged duration stimulation sometimes resulted in an additional, post-stimulus, haemodynamic response and a negative surround response which is analogous to the negative BOLD fMRI signal seen in previous studies by both our group and others (Boorman et al., 2015, Devor et al., 2007, Shmuel et al., 2006, Boorman et al., 2010). The polarity of the detected change in local neural activity evoked by SST-expressing interneurons is dependent on cortical depth, with increased MUA occurring at superficial depths and reduced MUA occurring deeper in the cortex. Meanwhile, NOS1-expressing interneurons evoke reliable changes in haemodynamics in the presence of minimal changes in local neural activity.

Taking first the NOS1-expressing interneurons, we found that specific optical stimulation was sufficient to elicit large localised increases in blood volume and saturation. This finding confirms the general idea that GABAergic neuron activity can produce increases in blood flow (Anenberg et al., 2015, Vazquez et al., 2018) and vessel diameter (Uhlirova et al., 2016), but extends current understanding by showing that these effects can be obtained by specifically stimulating the sub-class of NOS1-expressing interneurons. We are confident that the effects we report are a consequence of the action of the LED on the channelrhodopsin in the NOS1-expressing neurons in our transgenic animals because identical optical stimulation in wild-type controls was without effect. These findings provide additional support for the proposal that NOS1 interneurons play a critical role in the fundamental process of neurovascular coupling (Duchemin et al., 2012). The effects in the current study are likely to be mediated through the release of NO from these neurons, which is known to be a potent vasodilator (Furchgott and Zawadzki, 1980, Ignarro et al., 1987, Palmer et al., 1987).

However, perhaps the most significant finding with NOS1 interneuron stimulation was that the robust haemodynamic response occurred with only a minimal change in local MUA. Compared with the effects of mild sensory stimulation where a large MUA response was associated with a moderate haemodynamic response (Figures 2A, 4A, 5A, 6A), selective stimulation of NOS1 interneurons caused a larger haemodynamic response but with only a small change in MUA (Figures 2B, 4B, 5B, 6B). These findings would be consistent with a small population of optically activated NOS1 interneurons (∼2% of neurons in the light-activated area (Valtschanoff et al., 1993)) being responsible for the small change in MUA, yet having a dramatic effect on local haemodynamic activity. If correct, it would mean they would be in a position to play a critical role in the coupling of neural and haemodynamic activity.

While it is unlikely that NOS1 interneurons play an exclusive role in neurovascular coupling, these results have demonstrated for the first time that a particular subpopulation of cortical interneurons can evoke robust haemodynamic responses without being associated with the large increases in overall neural activity that would normally be expected. This suggests that any procedure or intervention that specifically targets these populations of cortical interneurons is likely to uncouple the neurovascular relationship upon which interpretations of BOLD signalling as an indirect proxy for neural activity depends.

We turn now to the case of SST-expressing interneurons. Our results show that a short 2s stimulation of SST interneurons also produced a large increase in blood volume and saturation in the activated area. With a longer duration stimulation period (16s), a longer latency negative haemodynamic response developed in adjacent unstimulated tissue (n=9/16 experiments; Figure 7), where the central positive haemodynamic response was surrounded by an inhibitory zone in adjacent tissue (Figures 5B, 7A). This inhibitory response was characterised by a marked reduction in blood volume and saturation. The temporal dynamics and magnitude of this response are similar to the negative BOLD surround region reported in previous studies by our group and others (Boorman et al., 2015, Shmuel et al., 2006). Although Uhlirova et al. (2016) have previously reported post-stimulus, NPY-mediated, vasoconstriction in response to optogenetic stimulation of VGAT-expressing interneurons (Uhlirova et al., 2016), the present study is the first demonstration of reduced blood volume (indicative of vessel constriction) occurring concurrently with stimulation of a single population of cortical interneurons.

To understand better the neural basis of the centre-surround pattern of haemodynamic responses elicited by stimulating SST interneurons we performed simultaneous 2D-OIS and multi-channel electrophysiology in the central stimulated region. Given that under ostensibly constant experimental conditions the surround haemodynamic inhibition was observed in some cases but not others, we sought to take advantage of this response variability by comparing the electrophysiological responses in the stimulated region when surround inhibition was present and when it was absent (Figure 7). In both instances MUA responses in the superficial cortical layers were similar. However, only in cases where the negative surround response was present, was there a strong suppression of MUA in the deep cortical layers. Then in the stimulation off-set period, after a transient increase in MUA activity, we observed a prolonged period (∼20s) of suppressed MUA below baseline across all layers.

A novel aspect of the present experiments with SST interneuron stimulation was the presence of a haemodynamic off-set response when the 16s stimulation was terminated. This effect was observed in some cases but not others. The reason for this is unclear, although slight variations in the depth of anaesthesia could be responsible. Thus, only in those cases where photostimulation induced a surround inhibition response was the prolonged offset response in the central, previously stimulated, region observed. If the central stimulation failed to produce surround inhibition then no positive offset response was observed. This later secondary peak in Hbt may be related to the long-lasting inhibition which is observed in these animals following photoactivation.

These responses during and after the 16s SST-stimulation period are difficult to interpret in terms of normal activation-induced neurovascular coupling.

Firstly, there seems to be a strong association between the presence of MUA inhibition in deep cortical layers and the inhibitory haemodynamic response in surrounding tissue. How might central activation of SST interneurons reduce blood volume in surrounding regions? Karnani et al. (2016) showed that lateral inhibition between adjacent cortical regions is mediated by lateral projecting SST interneurons. We have previously shown that a decrease in deep layer multi-unit activity is correlated with the negative surround BOLD signal in rat sensory cortex (Boorman et al., 2015, Boorman et al., 2010), our results presented here are qualitatively similar, suggesting SST interneurons projecting into neighbouring regions may be responsible. To our knowledge, this is the first demonstration of reduced haemodynamics resulting from the stimulation of a specific population of interneurons and, as such, furthers our understanding of neurovascular function relating to the activity of specific cell types.

Secondly, there was a similarly strong association between inhibition of deep-layer MUA and the occurrence of a large offset haemodynamic response in the previously stimulated region. It is particularly important to note that the positive post-stimulus haemodynamic response occurred without any corresponding increase in MUA. Indeed, the central area MUA was suppressed during this positive haemodynamic response. In light of our findings with the NOS1 interneurons (see above), one possibility could be that following SST interneuron stimulation offset, the small population of NOS1 cells in the central stimulated region become active, potentially due to removal of inhibition from the activated SST interneurons, thereby causing the large haemodynamic response during a period of overall MUA suppression (Figure 7C,E). Alternatively, Mariotti et al. (2018) reported that activation of cortical SST interneurons caused delayed long-lasting [Ca2+]_i_ elevations in astrocytes. Increased astrocyte [Ca2+]_i_ is known to produce significant vasodilation (Zonta et al., 2003, Gordon et al., 2008, Lind et al., 2013), therefore SST interneuron-evoked astrocyte [Ca2+]_i_ increases could cause the observed positive post-stimulus haemodynamic response. However, whether the time course of SST activation of astrocytes accords with the other neural and haemodynamic changes observed here remains to be determined.

While the use of anaesthesia in this study may be considered a limitation (Gao et al., 2017), we have previously demonstrated that in our chronic preparation (as used in the present study) mechanical somatosensory-evoked haemodynamic responses are comparable to those observed in the awake animal (Sharp et al., 2015).

Overall, the results of this study extend our knowledge of how specific subpopulations of cortical GABAergic interneurons mediate key aspects of neurovascular control. This has implications for our understanding of several diseases in which neurovascular coupling and inhibitory interneurons are dysfunctional or lost; including epilepsy (Dudek and Shao, 2003, Kumar and Buckmaster, 2006, Harris et al., 2013) and Alzheimer’s disease (Zlokovic, 2010, Verret et al., 2012). Indeed, the demonstration that the targeting of a single cell population reliably evokes robust haemodynamic changes in the absence of associated large increases in neural activity suggests a potential novel treatment strategy for diseases in which chronic hypoperfusion plays a role, such as Alzheimer’s disease. Meanwhile, the novel demonstration of deep layer inhibition during specific activation of SST interneurons could potentially be used to shut down aberrant neuronal activity, thus offering therapeutic strategies for diseases such as epilepsy. Future work should focus on the relationship between interneuron deficits, dysfunctional neurovascular coupling, and disease progression.

## Materials and Methods

### Animals

All animal procedures were performed in accordance with the guidelines and regulations of the UK Government, Animals (Scientific Procedures) Act 1986 and approved by the University of Sheffield Ethical review and licensing committee. Mice had ad libitum access to food and water and were housed on a 12 hour dark/light cycle. We used 39 mice of both sexes including 18 nNOS-CreER x ChR2-EYFP mice (M/F, 19-33g); 12 Sstm2.1Crezjh/j x ChR2-EYFP mice (M/F, 22-44g), 5 C57Bl/6J mice (F, 23-25g) and 4 non-ChR2-expressing littermates of NOS1-ChR2 mice (M, 33-39g). Sstm2.1Crezjh/j x ChR2-EYFP (SST-ChR2) mice were obtained by crossing homozygous SOM-IRES-Cre mice (Stock 013044, Jackson Laboratory, USA) with homozygous ChR2(H134R)-EYFP mice (Stock 024109, Jackson Laboratory), as described previously (Elghaba et al., 2016).

The nNOS-CreER x ChR2-EYFP (NOS1-ChR2) mice were obtained by crossing heterozygous nNOS-CreER (Stock 014541, Jackson Laboratory) with homozygous Ai32 mice (Stock 024109, Jackson Laboratory). Littermates lacking the nNOS-CreER insertion do not express ChR2 and were used as control mice in this study. ChR2 expression was induced by intraperitoneal (IP) injection of Tamoxifen (Sigma-Aldrich, Gillingham, UK) at 100 mg/kg, administered 3 times over 5 days. Treatment with tamoxifen was carried out a minimum of 2 weeks prior to surgery, to allow for gene expression to take place.

### Preparation of Chronic Cranial Window

A thinned cranial window was prepared over the right whisker barrel cortex, as previously described (Sharp et al., 2015). Anaesthesia was induced through intraperitoneal injection of fentanyl-fluanisone (Hypnorm, Vetapharm Ltd, Leeds, UK), midazolam (Hypnovel, Roche Ltd, Welwyn Garden City, UK) and sterile water (in a ratio of 1:1:2 by volume; 7ml/kg) and maintained using isoflurane (0.5-0.8%) in 100% oxygen at a flow rate of 1L/min. All surgeries were carried out in a dark room using a surgical illuminator with a band pass filter (577 ±5nm) to avoid erroneous optogenetic activation in the ChR2-expressing mice. Mice were placed on a stereotaxic frame (Kopf Instruments, Tujunga, USA), on a homeothermic blanket (Harvard Apparatus, Cambridge, UK) maintaining rectal temperature at 37°C. The bone overlying the right somatosensory cortex was thinned to translucency using a dental drill, forming an ≈3mm^2^ optical window. A thin layer of clear cyanoacrylate was applied and a stainless steel head plate secured to the skull using dental cement (Super bond C&B, Sun Medical). Surgery was performed at least 2 weeks before the first experimental imaging session.

For experiments, anaesthesia was induced as described above and maintained using isoflurane (0.25-0.7%) in 100% oxygen at a flow rate of 0.8L/min. Mice were placed on a stereotaxic frame and head fixed via their head plate. Animals were placed on a homeothermic blanket maintaining rectal temperature at 37°C.

### 2-Dimensional optical imaging spectroscopy (2D-OIS)

Animals underwent two experimental sessions. Session one, at least two weeks post-surgery, involved 2D-OIS recordings and application of both short and long duration whisker and light stimulations. Session two, occurring at least one week after session 1, involved concurrent electrophysiology and 2D-OIS recordings while applying both short and long duration stimulations (see below for full details).

As described previously (Berwick et al., 2005), 2D-OIS was used to record changes in cortical haemodynamics, allowing the estimation of changes in cortical oxyhaemoglobin (HbO_2_), deoxyhaemoglobin (Hbr) and total haemoglobin concentration (Hbt). The cortex was illuminated with 4 wavelengths of light (587±9 nm, 595±5nm, 560±15 nm, 575±5 nm) using a Lambda DG-4 high-speed filter changer (Sutter Instrument Company, USA). The re-emitted light was collected at a frame rate of 32Hz using a Dalso 1M60 CCD camera which was synchronized to the filter switching, thus producing an effective frame rate of 8 Hz. The camera was fitted with a 490 nm high pass filter to prevent light from the photostimulation LED being collected with the re-emitted light. The spatial maps recorded from the re-emitted light then underwent spectral analysis based upon the path length scaling algorithm (PLSA) described previously (Berwick et al., 2005, Mayhew et al., 1999). In brief, the algorithm uses a modified Beer-Lambert Law with a path length correction factor to convert detected attenuation in the re-emitted light into predicted absorption. These absorption values were then used to generate estimates of the changes in HbO_2_, Hbr and Hbt from baseline values. The concentration of haemoglobin in tissue was assumed to be 100 µM and oxygen saturation was assumed to be 70%. This spectral analysis produced 2D images of the micromolar changes in volume of HbO_2_, Hbr and Hbt over time.

### Stimulations

Whisker stimulation was performed using a plastic T-bar attached to a stepper motor, which deflected the whiskers ≈1cm in the rostro-caudal direction (Sharp et al., 2015). Whiskers were deflected at 5Hz for either 2 seconds or 16 seconds. To improve the signal-to-noise for each experiment whisker stimulation was presented 30 times in trials lasting 15-25 seconds each (2s stimulation) or presented 15 times in trials lasting 70 seconds each (16s stimulation).

Photostimulation was performed using a fibre-coupled LED light source (470m, ThorLabs, Newton, USA) and was delivered to the point of activation using a fibre optic (core diameter 200µm, ThorLabs). The cortex was illuminated for 2s (pulse width 10ms, 20Hz, 2V or 99Hz, 1V) or 16s (pulse width 10ms, 20Hz, 1.5V or 99Hz, 0.5V). These parameters were titrated to produce haemodynamic responses which were similar to physiologically-evoked responses. To improve the signal-to-noise for each experiment photostimulation was presented 20-40 times in trials lasting 15-25 seconds each (2s stimulation) or presented 15 times in trials lasting 70 seconds each (16s stimulation).

### Electrophysiology

In order to concurrently measure haemodynamic and neural responses, in a final imaging session (occurring at least one week subsequent to an imaging session in which only 2D-OIS data was acquired) a 16 channel microelectrode (100μm spacing, 1.5-2.7MΩ impedance, site area 177 µm^2^; NeuroNexus Technologies, Ann Arbor, USA) was inserted into the whisker barrel cortex through a small cranial burr hole. The electrode was positioned in the centre of the whisker region (area showing the largest blood volume response to a 2s mechanical whisker stimulation, defined by 2D-OIS in the previous imaging session) and inserted to a depth of 1500-1600µm. The electrode was connected to both a preamplifier and data acquisition device (Medusa BioAmp/RZ5, TDT, Alachua, USA). Data was sampled at 24kHz. For analysis, data was downsampled to 6kHz and multiunit activity (MUA) analysed.

### Analysis

Analysis was performed using MATLAB (MathWorks). Using the 2D-OIS-generated spatial map of HbO_2_ response to stimulation, a region of interest (ROI) was selected for use in subsequent time-series analysis. In order to select an ROI (red ROI in Figures 2, 5), the HbO_2_ image is processed to remove edge pixels and a Gaussian 3×3 filter applied. For the response period, each pixel is averaged across time to generate a mean value. The threshold for a pixel to be included in the ROI was set at 1.5 x standard deviation. Thus the ROI represents the area with the largest haemodynamic response to the stimuli. For each experimental paradigm, the response across all pixels in the ROI was averaged in order to generate a time series for each haemodynamic profile (Hbt, HbO_2_ and Hbr). As described above haemodynamic data were acquired with a 490nm high pass filter, resulting in minimal light artefact from the stimulating LED. For 20Hz stimulation this residual artefact was removed using a high pass filter followed by a modified boxcar function, for 99Hz stimulation only the modified boxcar function was applied. For each experiment, mean time series for Hbt, HbO_2_ and Hbr were produced by averaging across trials. Matching experiments (e.g. 2s photostimulation) from the same mouse on different imaging days were averaged together, so that each mouse only contributed one time series per experiment.

For the negative surround analysis, an automated ‘negative response’ ROI was selected, with a threshold of 1.5 x standard deviation, as described above. In mice in which no negative surround response was observed, ‘negative response’ ROIs were selected manually in an equivalent brain region.

### Statistical analysis and experimental design

In order to compare haemodynamic responses we focused on Hbt, which is our most reliable measure. Statistical comparisons were performed using GraphPad Prism (La Jolla, USA). A two-way ANOVA with Tukey’s multiple comparisons test was performed in order to undertake intergroup comparisons of MUA for different electrode depths in the presence and absence of a negative surround haemodynamic response. A one-way ANOVA with Tukey’s multiple comparisons test was used to compare Hbt peaks in response to various paradigms (whisker stimulation, photostimulation of NOS-ChR2 mice, photostimulation of SST-ChR2 mice), and to compare photostimulation-evoked MUA in different mouse lines (NOS-ChR2 mice, SST-ChR2 mice, control mice). T-tests (paired or unpaired, as appropriate) were used to compare between two groups (MUA in superficial vs deep electrode channels and initial vs secondary peak haemodynamic response). *P* values <0.05 were considered to be statistically significant. Data are presented as mean ± standard error in the mean (s.e.m). Experiments and analysis were performed unblinded to mouse type.

### Immunohistochemistry

At the end of the experiments, mice were given an overdose of pentobarbital and perfused with saline followed by 4% paraformaldehyde (PFA) in phosphate-buffered saline (PBS), via cardiac perfusion. Brains were extracted, postfixed in 4% PFA for 24 hours at 4°C and cryoprotected in 30% sucrose in PBS at 4°C for 48 hours. The brains were sectioned using a cryostat (Thermo Fisher Scientific, Loughborough, UK) to produce 30 µm coronal sections which were placed into PBS with 0.3% Triton X-100 (PBST) for 20 minutes to permeablize the tissue. Free-floating sections were blocked with 10% normal donkey serum (Jackson ImmunoResearch, Ely, UK, when staining for NOS1 and EYFP) or 10% normal rabbit serum (Vector Laboratories, Peterborough, UK, when staining for SST) in PBST for 1 hour, and then incubated overnight at 4°C in blocking solution with primary antibodies against GFP (anti-GFP, rabbit polyclonal, [ab290], Abcam, Cambridge, UK, 1:200, recognises EYFP, which is a genetic mutant of GFP); NOS1 (anti-nNOS, goat polyclonal, [ab1376], Abcam, 1:250) or SST (anti-Somatostatin, rat monoclonal (YC7), [MAB354], Millipore, Watford, UK, 1:500). Sections were washed with PBS and incubated in the appropriate fluorescent secondary antibodies: Alexa 568 donkey anti-rabbit (Thermo Fisher Scientific, 1:500 dilution in PBS), Alexa 488 donkey anti-goat (Thermo Fisher Scientific, 1:500 dilution in PBS) for 2 hours at room temperature, or in the case of the SST stain: incubated in biotinylated rabbit anti-rat (Vector Laboratories, 1:100 dilution in PBS with 1.5% normal rabbit serum) for 30mins at room temperature. The SST staining sections were then washed with PBS and incubated with Alexa 647 Streptavidin (Thermo Fisher Scientific, 1:400 dilution with PBS) for 90 minutes. All sections were mounted onto gelatin coated slides following immunohistochemical staining and imaged with a fluorescence stereo microscope (M205 FA, Leica Microsystems, Milton Keynes, UK).

## Acknowledgements

We would like to thank Michael Port for building and maintaining the whisker stimulation device and 2D-OIS apparatus.

